# Modeling microbial communities using biochemical resource allocation analysis

**DOI:** 10.1101/537779

**Authors:** Suraj Sharma, Ralf Steuer

## Abstract

To understand the functioning and dynamics of microbial communities remains a fundamental challenge at the forefront of current biology. To tackle this challenge, the construction of computational models of interacting microbes is an indispensable tool. Currently, however, there is a large chasm between ecologically-motivated descriptions of microbial growth used in ecosystems simulations, and detailed metabolic pathway and genome-based descriptions developed within systems and synthetic biology. Here, we seek to demonstrate how current biochemical resource allocation models of microbial growth offer the potential to advance ecosystem simulations and their parameterization. In particular, recent work on quantitative microbial growth and cellular resource allocation allow us to formulate mechanistic models of microbial growth that are physiologically meaningful while remaining computationally tractable. Biochemical resource allocation models go beyond Michaelis-Menten and Monod-type growth models, and allow to account for emergent properties that underlie the remarkable plasticity of microbial growth. We exemplify our approach using a coarse-grained model of cyanobacterial phototrophic growth, and demonstrate how the model allows us to represent physiological acclimation to different environments, co-limitation of growth by several nutrients, as well as emergent switches between alternative nutrient sources. Our approach has implications for building models of microbial communities to understand their interactions, dynamics and response to environmental changes.

## INTRODUCTION

Microbial organisms and their metabolism are integral parts of the Earth’s biogeochemical cycles, and play key roles in almost all ecosystems and environments. Microbial organisms typically form complex, interacting and dynamically changing communities, with examples ranging from the gut microbiome to marine plankton communities. To understand the organizing principles and the functioning of such communities is of paramount importance for a vast number of basic and applied research questions, including questions pertinent to biotechnology, climate change, and human health [28, 61, 21, 56]. Despite the significant advances in our ability to observe and characterize biological systems, however, understanding the interactions and the emergent dynamics of microbial communities remains a fundamental, and truly transdisciplinary, challenge [15, 1, 30, 65, 61].

Traditionally, ecosystem dynamics and microbial communities are the realm of microbial ecology, with a long history and a wealth of results concerning the organization, stability, and functioning of (microbial) ecosystems [28, 58]. In the past century, a variety of modeling approaches have been developed to address fundamental ecological questions, ranging from understanding patterns of biodiversity to predicting the response of ecosystems to changing environmental conditions [16, 15, 30, 58]. Descriptions of microbial growth range from phenomenological ‘black box’ models to more elaborate trait-based models of growth [15, 30]. It has been noted though, that current theoretical approaches to microbial growth are still dominated by the classic Monod or Michaelis-Menten functional form [1, 22]. While undoubtedly highly successful, Monod-type models of growth exhibit a number of limitations. For example, it has been argued that the constant parameters used in the Monod equation cannot account for the observed plasticity of microbial physiology [5]. Likewise, it has been noted that, despite the significant advances in genome sequencing and quantitative high-throughout methods, the complexity of mechanistic ecosystem models, and in particular the description of microbial growth within these, have not changed substantially since they were developed in the 1970s [22].

Parallel to progress in theoretical and experimental microbial ecology, the past two decades have seen an unprecedented advance in our understanding of microbial molecular physiology—mainly driven by advances in our ability to monitor, measure and modify the inner workings of cells. Theoretical and computational descriptions of microbial metabolism, facilitated by large-scale metabolic network reconstruction and constraint-based analysis, have become established tools in molecular systems biology [45, 60, 8]. Curated genome-scale reconstructions of metabolic networks are available for an increasing number of microbial organisms [23, 2, 32], and are increasingly recognized as a resource in studies of microbial communities [64, 56, 21, 65, 61, 33],

More recently, also the quantitative physiology of bacterial growth has gained renewed attention, with numerous studies providing insights into the principles of microbial growth and resource allocation [41, 53, 52, 11, 63]. Key observations concern the ‘laws’ and trade-offs of microbial growth. In continuation of the classic studies of bacterial growth physiology, a number of studies have recently addressed the covariation between the cellular composition and the growth rate of microorganisms [53, 52]. Theoretical descriptions of microbial resource allocation include coarse-grained models that describe the fundamental processes of cellular growth by partitioning the proteome into few essential classes [38, 35, 59, 50, 11], as well as large-scale constraint-based models that take into account the costs and benefits of each individual gene [18, 42, 19, 49]. In particular, the concept of resource balance analysis [18, 19] and related methods, such as such as metabolism and macromolecular expression models [42], show that quantitative models that predict protein expression and the cellular composition are feasible on the genome-scale—and can be extended to time-varying environments [49]. As yet, however, quantitative modeling of microbial resource allocation is mostly restricted to well characterized model organisms in typical laboratory or biotechnological environments.

The purpose of this work is therefore to outline a bridge between these recent studies on microbial resource allocation and current models of microbial ecology. We argue that biochemical resource allocation models offer significant potential to advance ecosystem simulations beyond current applications of constraint-based analysis of microbial metabolism. In particular, we seek to demonstrate that biochemical resource allocation models, as defined below, can be constructed and parameterized for large classes of microbial organisms based on available biochemical and physiological data; and are, unlike Monod-type models, capable to exhibit emergent properties of growth, such as switching between alternative sources of nutrients. Our study is motivated by recent calls for a new generation of plankton models to better capture the emergent properties of marine ecosystems [1, 22]. As will be demonstrated below, biochemical resource allocation models follow the rationale described by Allen and Polimene [1] to design a generic cell model that captures the essence of key physiological activities and that is based on a robust physiological formulation of competing physiological activities — and therefore should be able to reproduce biogeochemical and ecological dynamics as emergent properties.

The paper is organized as follows: In the first section, we briefly recapitulate computational models of microbial growth. In the second section, we provide an overview of metabolic network reconstruction and recent biochemical models of cellular resource allocation. In the following section, we consider a coarse-grained model of phototrophic (cyanobacterial) growth and describe its parameterization. In the subsequent section, we then discuss emergent properties of the model, in particular cellular growth laws and co-limitation, as well as the representation of microbial diversity and the uptake of alternative nutrients. In the sixth and seventh sections, we present a brief case study: the co-existence of two phytoplankton species with a gleaner-opportunist trade-off. In the final section, we provide a discussion and outlook.

## MODELS OF MICROBIAL GROWTH

Assuming a chemostat-like setting, the growth dynamics of a population of (genetically homogeneous and well mixed) cells can be described by an ordinary differential equation,

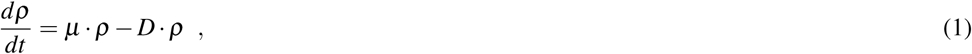

where *ρ* denotes the concentration of cells (in units of cells per volume), *µ* denotes the specific growth rate and *D* is the dilution or death rate. The specific growth rate *µ* is a function of the respective environment, and depends on the concentrations of one or more nutrients. Typically, a limiting nutrient *n* with concentration [*n*] is considered that is supplied with the inflow of fresh medium at a concentration [*n*_*x*_]. The respective nutrient dynamics are described by

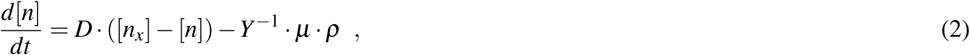

where *Y* denotes the yield coefficient, defined as the number of microbial cells (or units biomass) per unit of nutrient. The model can be readily extended to multiple microbial strains with concentrations *ρ*_*i*_ and several nutrients *n*_*k*_. The dynamics of a chemostat have been studied extensively [58] and equations of the form (1) and (2), as well as the respective extension to multiple strains and nutrients, are commonly utilized in ecosystem simulations [16, 15, 58, 57].

To evaluate the dynamics of the system requires knowledge of the specific growth rate *µ* as a function of the concentration of the limiting nutrient *n*. To this end, the most widely used approach is still to make use of the hyperbolic dependency proposed by Jacques Monod in 1949 [39],

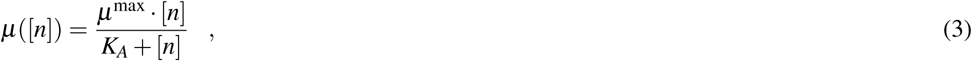

where *µ*^max^denotes the maximum specific growth rate of the microorganism in this environment and *K*_*A*_ denotes the half-saturation constant. The Monod equation is identical to the Michaelis-Menten equation of enzyme kinetics and represents an empirical description of microbial growth. Its constant parameters are typically estimated for specific environmental conditions and reflect a particular strain or species and its functional traits related to nutrient uptake and growth [5, 30]. Over the past decades, there have been several advances and alternative formulations of growth models, such as the Droop model [9] that introduces internal nutrient quotas. For phototrophic microorganisms, modifications are typically required to account for the effects of photoinhibition—the decrease of the specific growth rate for high light intensities [11]. A widely equation for phototrophic growth in dependence of the light intensity *I* is the Haldane equation,

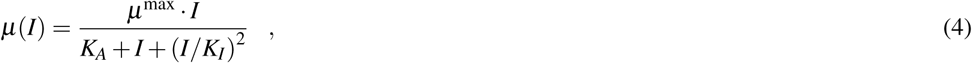

where *K*_*I*_ denotes the impact of photoinhibition. In the absence of photoinhibition (*K*_*I*_ →∞) the model is identical to the Monod equation with light as the limiting substrate. See, for example, Lee et al. [29] for a review on empirical growth models and their parameterization for different microalgae. The use of the Monod and related equations remain ubiquitous in current models of ecosystems [16, 5, 12, 30, 22, 57]. It has been emphasized recently [1, 22], however, that empirical growth models do not necessarily reflect our vast recent increase in knowledge about the quantitative physiology of microbial growth. The challenge before us is therefore to combine the conceptual simplicity of empirical growth models with molecular properties of microbial growth.

## METABOLIC RECONSTRUCTIONS AND CELLULAR RESOURCE ALLOCATION

Models of microbial and phytoplankton growth that incorporate internal structure and aspects of physiology are not new. Examples include the (still empirical) model of Droop [9] as well as other ‘internal-quota’ models—each representing a cell with one or more internal variables, and typically allowing for adjustments in the composition of cellular biomass [15]. Likewise, models that incorporate cost-benefit consideration have been proposed, most notably by JA Raven [48] and RJ Geider [17]. In the following, we build on these ideas and incorporate recent approaches to biochemical models of microbial growth [8, 60].

In particular, over the past two decades, genome-scale reconstructions (GMRs) of microbial metabolism have reached maturity and are available for a rapidly increasing number of (sequenced) microbial organisms. GMRs provide a comprehensive account of biochemical interconversions between small molecules (metabolites) within a cell or organism, and therefore allow to accurately estimate the stoichiometric and energetic synthesis costs of cellular constituents. GMRs have been highly successful to predict maximal growth yields of microbial organisms and other proerties of biotechnological relevance [45]. More recently, large-scale constraint-based resource allocation models [18, 42, 19, 49] were introduced that allow to predict protein expression and cell compositions of microbes in specified (albeit, with the exception of [50, 49], constant) environments. These models are based on the insight that the (maximal) flux of an enzyme-catalyzed biochemical reaction is typically constraint by the amount of the respective enzyme. Since enzymes are itself the products of metabolism, incorporating enzyme-dependent flux constraints gives rise to a self-consistent description of microbial growth: for any given growth rate *µ* the set of cellular enzymes must be sufficient to sustain the synthesis of the required precursors to allow for the translation of the set of catalyzing enzymes itself, as well as for the synthesis of all other (non-enzymatic) compounds within a cell.

More formally, the synthesis rate of a cellular protein *P*_*k*_ can be described by the equation

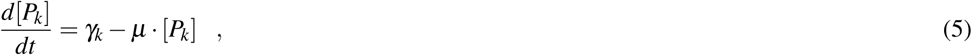

where *γ*_*k*_ denotes the translation rate of the protein that is required to match the dilution term *µ* [*P*_*k*_] of cellular growth (protein degradation can be readily included but is neglected in the following). The sum of all translation rates is constrained by the available ribosomal capacity and hence by the number of ribosomes.

To account for the synthesis of metabolic precursors and other cellular components, the interconversion of internal metabolites *m* is described by a stoichiometric matrix *N* and a vector *ν* that denotes the rates of (spontaneous or enzyme-catalyzed) interconversion rates,

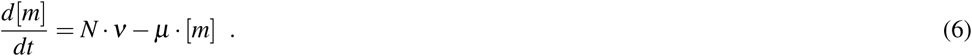

Typically, intracellular metabolism is assumed to be at steady state and the dilution terms for intracellular metabolites are neglected due to the high turnover of metabolites compared to their dilution by growth. In this case, the mass-balance constraint on intracellular reaction fluxes simplifies to *N · v* = 0.

To account for biochemical resource allocation, the rates of those reactions that are catalyzed by proteins are constrained by the amount of the respective catalyzing proteins

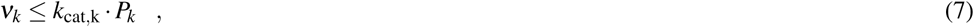

where *k*_cat,k_ denotes the specific activity of the enzyme or protein. The maximal uptake rate *ν*_*T*_ of an external nutrient *n*_*x*_ can be further constrained by the concentration of the respective nutrient and the amount of the respective transporter complex *P*_*T*_.

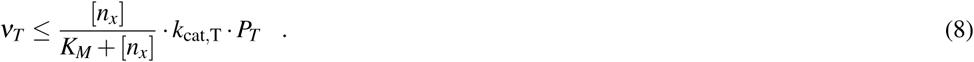

The uptake constraints can be modified to, for example, also account for diffusion limitations of nutrient uptake described by Bonachela et al. [5]. The constraints and equations summarized above, together with the assumption of a constant cell density, provide a quantitative description of microbial growth that is based on linear constraints. To obtain an estimate of the growth rate for a specific environment, the model is solved using the assumption that, during evolution, the fluxes are organized such that they give rise to a maximal growth rate in the respective environment (assumption of evolutionary optimality). Hence, similar to flux-balance analysis [45] and other constraint-based analysis, the assumption of optimality replaces unknown regulatory mechanisms.

The required parameters for model construction are: (i) the metabolic network (as encoded in the stoichiometric matrix *N* and the associated enzyme-reaction relationships). These data are available as part of a metabolic network reconstruction; (ii) the composition of the catalyzing enzymes (in terms of amino acids and possible co-factors). For most enzymes this information is readily available and part of reaction databases; as well (iii) as the specific activity *k*_cat_ of each catalyzing enzyme and, if required, the half-saturation constants for transporter reactions. While quantitative data is still scarce, in particular for non-model organisms, specific activities for a wide range of enzymes can be sourced from suitable databases, such as BRENDA [26], and are therefore (at least approximately) available. As we have argued previously [50, 49], reasonable estimates for all required parameters exist.

We note that the respective models can be constructed either on a genome-scale, involving all known individual enzymatic reaction steps of the respective organisms, see for example Goelzer et al. [19] or Reimers et al. [49]. Or, alternatively, by defining coarse-grained enzyme complexes that represent classes of reactions or pathways, see for example Rügen et al. [50]. Computationally, for any given growth rate, the resource allocation model gives rise to a linear program (LP) and hence can be solved efficiently. The maximal growth rate is then identified using bisection, see Materials and Methods for computational details. In the following, we refer to the type of models outlined above as biochemical resource allocation models (BRAMs).

## A MODEL OF PHOTOTROPHIC GROWTH

To exemplify the utility of biochemical resource allocation models for microbial ecology, we consider the construction and analysis of a coarse-grained model of cyanobacterial growth, based on previous work [50, 49, 11]. The model is depicted in Figure 1 and consists of a light harvesting reaction, 5 metabolic reactions involving 6 internal metabolites, as well as 8 catalyzing protein complexes and their respective translation reactions. In brief, inorganic carbon (*C*_*x*_) is taken up using an energy-dependent transporter (T_C_). Intracellular inorganic carbon is assimilated into organic carbon (*C*_3_) using inorganic carbon concentrating mechanisms and the Calvin-Benson cycle (CB). The metabolic intermediate *C*_3_ is converted into amino acids (AA) by a coarse-grained metabolism (M_C_).

**Figure 1.**
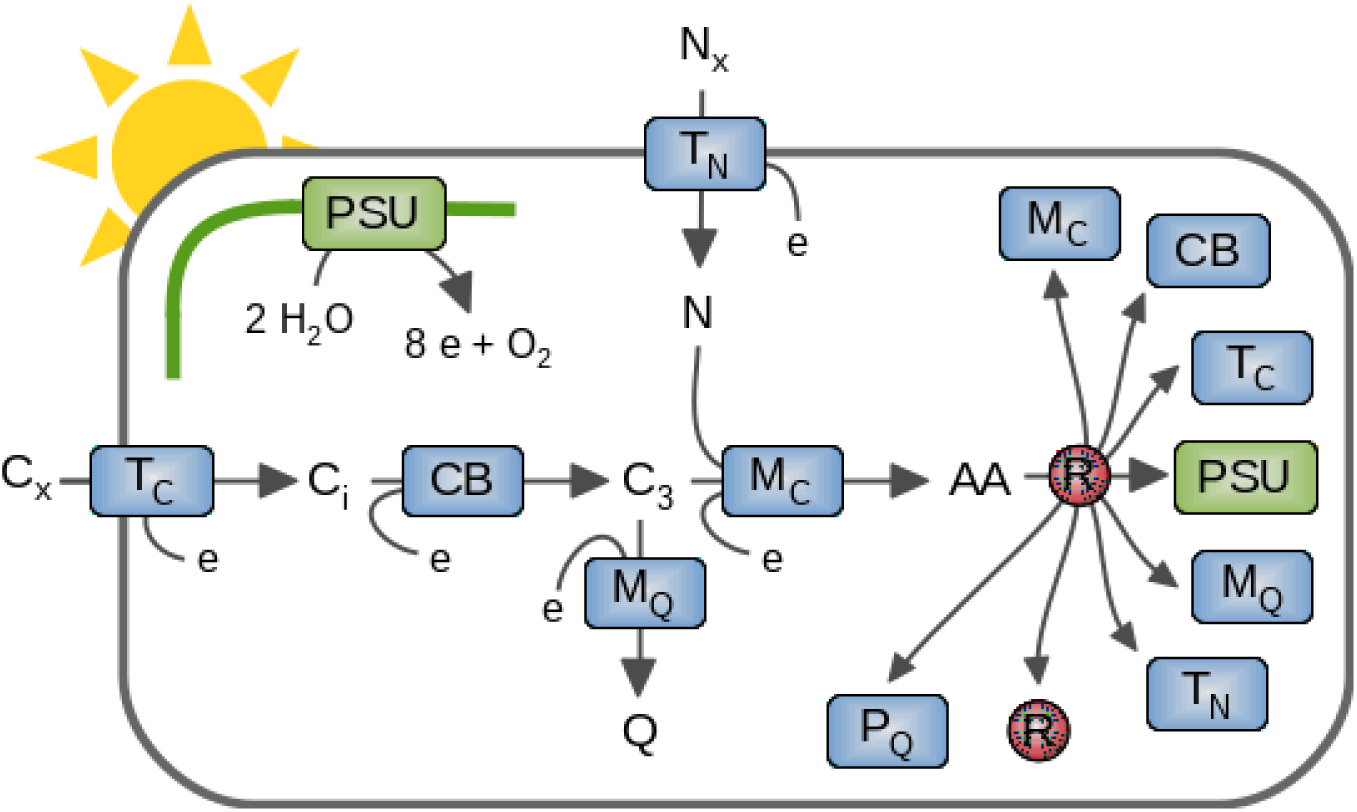
A coarse-grained biochemical resource allocation model of phototropic growth. The model consists of 8 protein complexes that catalyze 5 metabolic and transport reactions, as well as light harvesting and photosynthetic electron transport. Extracellular carbon (*C*_*x*_) is takes up and converted in amino acid (AA) precursors for translation of protein complexes by ribosomes (R). The abundances of protein complexes constrain cellular reaction rates. Abbreviations of protein complexes: photosynthetic unit (PSU), carbon uptake (T_*C*_), carbon assimilation (CB), nitrogen uptake and metabolism (T_*N*_), central metabolism and amino acid synthesis (M_*C*_), synthesis of other cellular constituents (M_*Q*_), and ribosomes (R). Abbreviations of metabolites: external inorganic carbon (C_*x*_), internal inorganic carbon (C_*i*_), assimilated organic carbon and precursor for biosynthesis (C_3_), amino acids (AA), external nitrogen (N_*x*_), internal nitrogen (N), remaining cellular constituents (Q), cellular energy unit (*e*).

The biosynthesis of amino acids requires a source of nitrogen (N) that is taken up from the environment using an energy-dependent transporter and associated nitrogen metabolism (T_N_). For amino acid synthesis, we assume a N:C ratio of ∼1*/*3 (the cellular N:C ratio is lower due to the remaining non-protein biomass component *Q*). Light harvesting and the photosynthetic electron transport chain are represented by a coarse-grained photosynthetic unit (PSU). The PSU protein complex regenerates cellular energy units *e* (combining ATP and reductant NADPH into a single energy unit). Amino acids are translated into proteins by ribosomes (R), which are itself protein complexes. The fraction of non-enzymatic proteins is represented by a (quota) protein component *P*_*Q*_. The remaining biomass is lumped into a metabolic component *Q* that is synthesized from the cellular precursor *C*_3_ by the protein complex M_Q_. All proteins complexes represent aggregates of individual proteins. The model assumes a constant cellular density. The specific growth rate is not dependent on cell size, but cell size may constraints parameters, such as the surface to volume ratio. The full set of equations is provided in the Materials and Methods.

All enzyme-catalyzed reactions are constrained by the amount of the respective enzymes. For example, carbon uptake is constrained by the equation

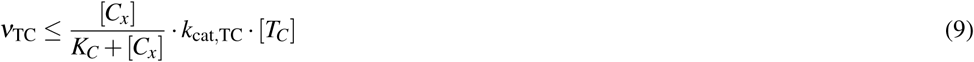

where [*T*_*C*_] denotes the amount of uptake transporter (in molecules per cell), *k*_cat,TC_ denotes the specific catalytic activity of the transport, and *K*_*C*_ denotes the half-saturation constant of the uptake complex. We note that Equation (9) provides an upper limit only, the actual flux can be less (for example by inactivating a fraction of the uptake transporter). Likewise, additional constraints can be included, such as an upper limit on the uptake flux induced by diffusion limitations [5]. Other intracellular reactions are constrained by the upper limits induced by the amount of the respective enzymes, e.g., for the carbon assimilation reaction

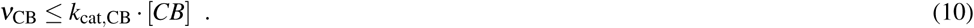

The model is parameterized using information about the individual enzymatic and biochemical processes. Using data from Faizi et al. [11], the effective size of the (coarse-grained) protein complexes can be approximated by the number of enzymes involved in amino acid synthesis multiplied with the average size (in units of amino acids) per enzyme. Catalytic turnover numbers *k*_cat_ are assigned according to typical values for the respective reactions. For example, the rate of translation per ribosome is approximately 20 amino acids per second, the photosynthetic unit (with photosystems II as rate limiting complex) is assumed to give rise to approximately 250 interconversion per second, the *k*_cat,TC_ of the carbon transporter is set to 20*s*^*-*1^, the catalytic activity of the central metabolism is set to *k*_cat,MC_ is set to 10*s*^*-*1^. Reasonable parameter ranges for many enzymatic processes (for a generic cell) can be obtained, for example, from Milo and Phillips [37]. The full set of parameters used in the following is provided in the Materials and Methods.

Given the stoichiometric constraints and the assigned parameters, the model gives rise to a global optimization problem, and solved as a series of LP problems to identify the maximal specific growth rate *µ* in dependence of the availability of extracellular nitrogen and carbon and light intensity *I* (assumption of evolutionary optimality). Figure 2 shows the resulting growth curves as a function of environmental parameters. Similar to previous models [11], the resulting growth curves with respect to external nitrogen (*N*_*x*_) and carbon (*C*_*x*_) concentrations are consistent with Monod kinetics, the dependence of the specific growth rate on the light intensity is consistent with the Haldane equation.

**Figure 2.**
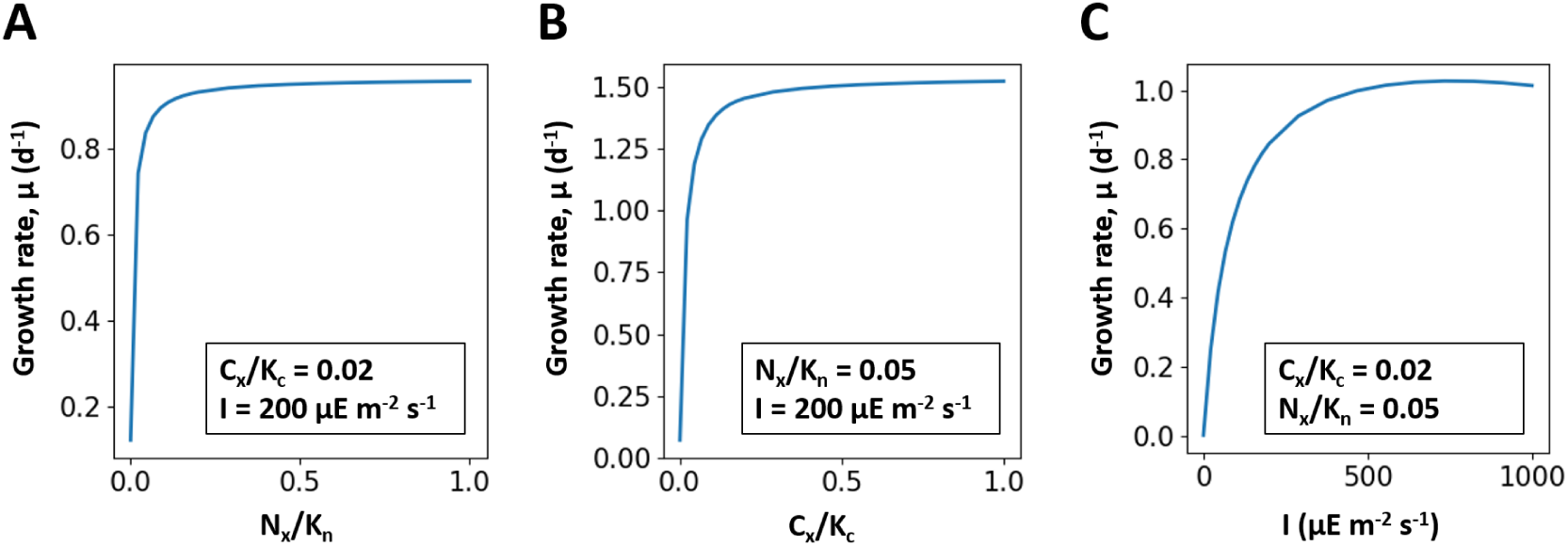
The maximal specific growth rate *µ* as a function of extracellular nutrient concentrations (*N*_*x*_ and *C*_*x*_) and light intensities *I*. Nutrient concentrations are reported relative to the half-saturation constant of the respective transporter complex. **Panel A:** The specific growth rate *ν*(*N*_*x*_*/K*_*n*_) with fixed *I* = 200*µEm*^*-*2^*s*^*-*1^and *C*_*x*_*/K*_*c*_ = 0.02. **Panel B:** The specific growth rate *ν*(*C*_*x*_*/K*_*c*_) with fixed *N*_*x*_*/K*_*n*_ = 0.05 and light intensity *I* = 200*µEm*^*-*2^*s*^*-*1^. **Panel C:** The specific growth rate *ν*(*I*) with fixed *C*_*x*_*/K*_*c*_ = 0.02 and *N*_*x*_*/K*_*n*_ = 0.05. Abbreviations: *N*_*x*_, external nitrogen concentration; *K*_*n*_, half-saturation constant of nitrogen transporter; *C*_*x*_, external inorganic carbon concentration; and *K*_*c*_, half-saturation constant of carbon transporter.

We emphasize that the growth curves shown in Figure 2 are emergent properties of the underlying constraints and parameters—and that changes in these constraints and parameters entail (sometimes complex) changes in overall growth properties. For example, the apparent half-saturation constant of the organismal growth curve is markedly different from the half-saturation constant assigned to the respective transporter complex, due to the fact that the cells can acclimate to low nutrient conditions by changing the expression of the respective protein complex.

## ACCLIMATION, TRADE-OFFS AND CO-LIMITATION

Biochemical resource allocation models go beyond describing nutrient uptake and the specific growth rate, and allow us to obtain insights into acclimation, co-limitation and cellular trade-offs. In particular, concomitant to the cellular growth curves, we obtain the distribution of protein resources within the cell as a function of environmental parameters and growth rate.

Figure 3 and Table 1 show the relative protein fractions invested into the different biochemical processes dependent on environmental conditions. The resource allocation framework allows the model to acclimate to the respective environmental condition and invest cellular resources into processes that would otherwise limit growth. As a consequence, the maximal uptake rate of the nitrogen transporter complex (*V*_max_ = *k*_cat,TN_ [*T*_*N*_]) and hence the affinity *A* = *V*_max_*/K*_*N*_ for the extracellular nitrogen source is not constant, but increases with decreasing external concentrations. Figure 4A shows the maximal uptake rate *V*_max_, as well as the actual uptake flux, as a function of extracellular nitrogen. Similar to the analysis by Bonachela et al. [5], and unlike descriptions using the Monod equation, the model accounts for the acclimation of the cell to low nutrient availability—with important consequences for, e.g., estimations of phytoplankton abundances in global ocean models. Likewise, protein investments in light harvesting strongly depend on the light intensity, at the expense of investments in other metabolic processes (Table 1).

**Table 1.** Cellular protein allocation in dependence of environmental conditions. Values denote the relative abundance (% relative to total proteome) of protein complexes under low and high nutrient conditions. The symbols hold same meaning as Figure 1.

| Protein | Low |  |  | High |  |  |
| --- | --- | --- | --- | --- | --- | --- |
|  | Nitrogen | Carbon | Light | Nitrogen | Carbon | Light |
| $T_C$ | 26.5 | 25.4 | 9.0 | 34.2 | 1.8 | 34.8 |
| $T_N$ | 15.2 | 9.4 | 2.4 | 0.9 | 14.8 | 9.4 |
| PSU | 23.2 | 30.2 | 53.5 | 29.9 | 47.8 | 20.8 |
| R | 5.0 | 5.0 | 5.0 | 5.0 | 5.0 | 5.0 |
| $P_Q$ | 20.0 | 20.0 | 20.0 | 20.0 | 20.0 | 20.0 |
| CB | 8.3 | 7.8 | 9.4 | 7.8 | 7.1 | 7.8 |
| $M_C + M_Q$ | 1.7 | 2.2 | 0.6 | 2.2 | 3.5 | 2.2 |

**Figure 3.**
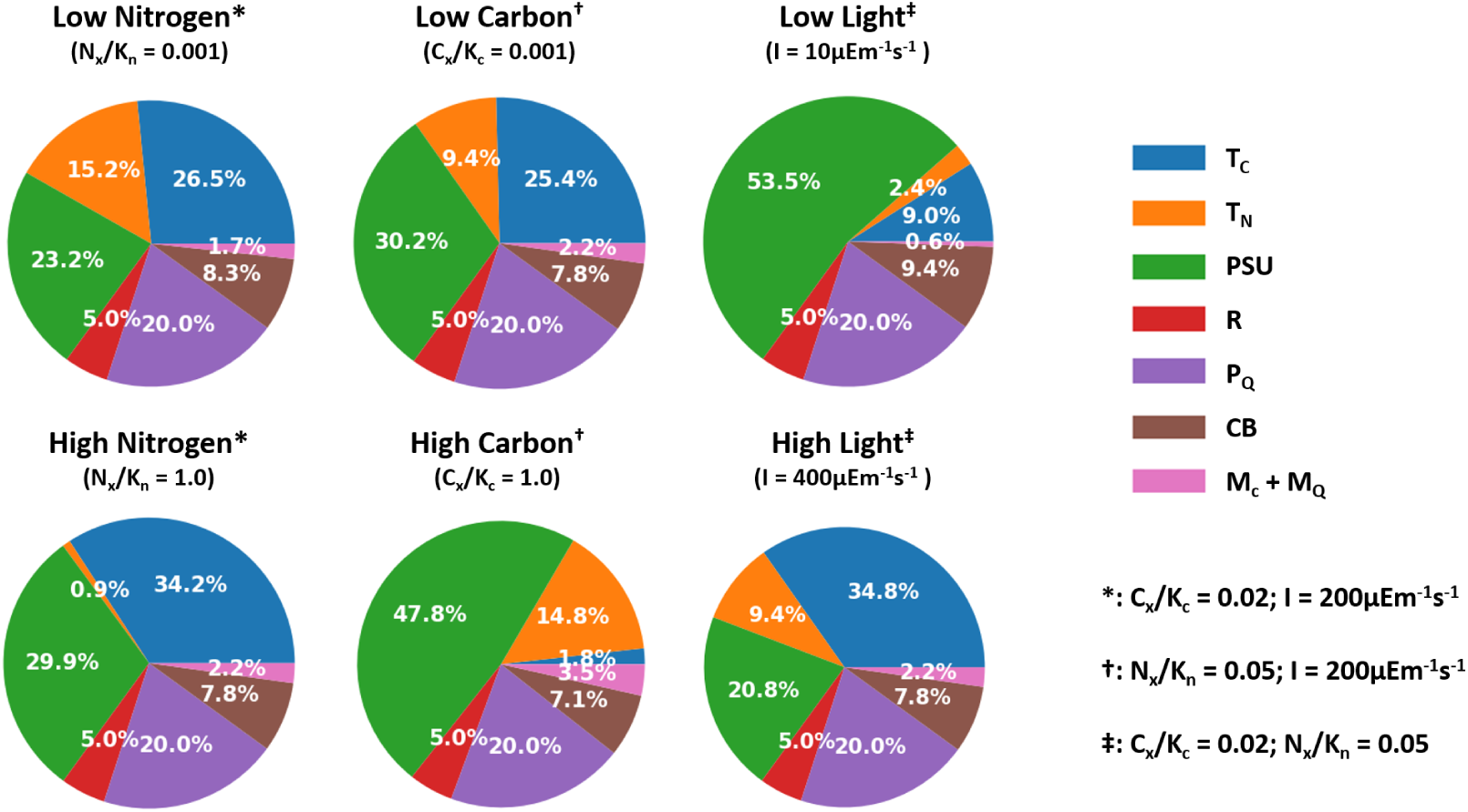
Cellular protein allocation in dependence of environmental conditions. Shown are the relative abundances of ribosomes and coarse-grained protein complexes under different growth conditions (relative to total protein but excluding the constant protein fraction *T*_*Q*_). The superscripts (∗, † and ‡) indicate the parameter values used to specify environmental conditions.

**Figure 4.**
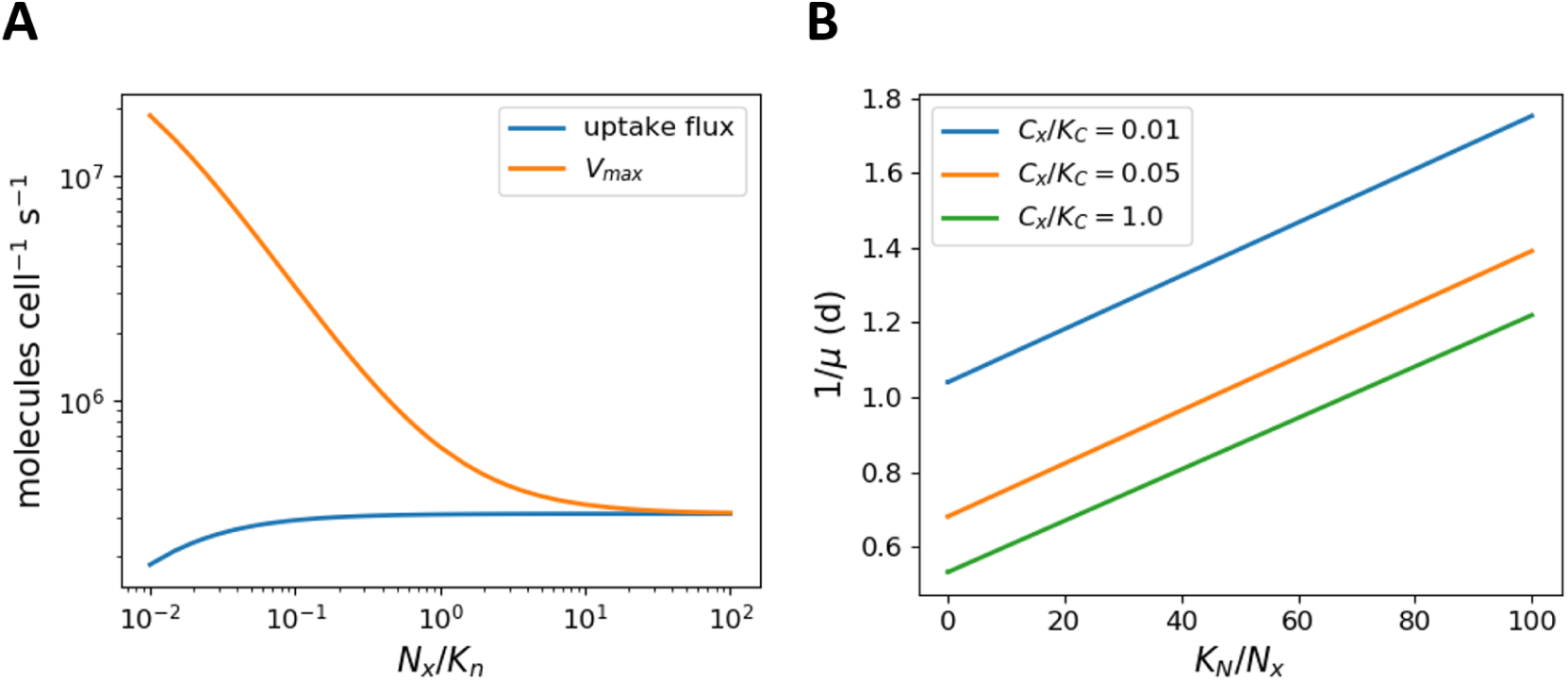
Panel A: The maximal uptake capacity *V*_max_ of the nitrogen transport complex (*V*_max_ = *k*_cat,TN_ [*T*_*N*_]) versus the actual uptake rate as a function of external nitrogen. For scarce nutrients more cellular resources are invested into the uptake capacity. **Panel B:** A Lineweaver-Burk plot of the (inverse of the) growth rate versus the (inverse of the) relative substrate concentration, *K*_*N*_ */N*_*x*_, for different values of external inorganic carbon. Parallel lines in a Lineweaver-Burk plot correspond to uncompetitive inhibition, whereas a multiplicative dependence of the growth rate on its substrates would result in lines with a identical x-intercept.

Closely related to trade-offs in resource allocation, it is an important challenge for empirical growth models to describe the dependence of growth on several potentially limiting nutrients, see Saito et al. [51] for a discussion on the concept and types of co-limitation. The most common ways to implement multiple limitation scenarios relies on either Liebig’s law of the minimum,

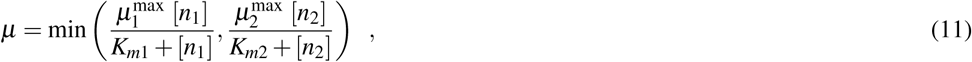

or the multiplicative form

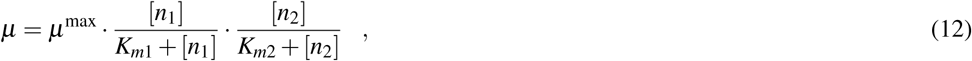

where [*n*_1_] and [*n*_2_] denote the concentrations of two potentially limiting nutrients and *K*_*m*1_ *K*_*m*2_ the respective half-saturation constants, respectively. As discussed by Saito et al. [51] both forms are not without problems and there is no clear empirical evidence to assess the merits of either representation. Given its simplicity, the multiplicative form is commonly employed in multi-nutrient models [16, 57].

For biochemical resource allocation models, however, the description of growth limitations as a function of two or more nutrients emerges without further assumptions about the functional form of growth equations. In particular, the coarse-grained model described above is not consistent with Liebig’s law of the minimum, as growth on a single nutrient, as shown in Figure 2, does not exhibit any hard threshold. The absence of such a threshold is due to the fact that, for scarce nutrients, resources are increasingly invested into the respective uptake reactions.

More relevant, however, the emergent growth curve is also not consistent with a multiplicative functional form. Figure 4B shows a Lineweaver-Burk plot of the growth rate as a function of nitrogen availability for different values of the external carbon concentration. Parallel lines in a Lineweaver-Burk plot correspond to uncompetitive inhibition, whereas a multiplicative functional form would result in lines with an identical x-intercepts. Hence, the absence of carbon acts analogous to uncompetitive inhibition, and affects both, the apparent organismal half-saturation constant of growth, as well as the maximal growth rate of the cell—again with important consequences for, e.g., growth limitations and nutrient dynamics in coupled ecosystem models.

## METABOLIC DIVERSITY AND THE COST OF REGULATION

Recent studies have emphasized microbial community diversity as a fundamental property of model ecosystems [15]. Several principles and mechanisms allow to represent microbial diversity in biochemical resource allocation models. A shown above, cells may acclimate to different environmental conditions, resulting in an inhomogeneous population. If required, the possibility for physiological acclimation can also be restricted within simulations, for example by allowing only limited ranges for intracellular protein complexes (as would be expected, for example, for cyanobacterial *Prochlorococcus* strains).

More importantly, however, cellular diversity also arises due to genetically-encoded differences between organisms. Firstly, diversity may arise due to differences in enzyme-kinetic parameters. The evolution of enzyme-kinetic parameters is constrained by physicochemical limits that result in trade-offs between parameters, with the protein complex ribulose-1,5-bisphosphate carboxylase/oxygenase (RuBisCO) as a prominent example [14]. As will be shown below, these differences result in different cellular growth curves. Secondly, microbial organisms exhibit metabolic diversity with respect to the encoded metabolic functionality within their genomes. As shown by recent studies of the cyanobacterial pan-genome and pan-metabolism [3, 55, 4], genome sizes differ significantly—reflecting the different adaptations and lifestyles of organisms. Differences in the set of encoded proteins give rise to different metabolic strategies that are accessible to the organism, for example with respect to the modes of energy generation [13], or accessibility of nutrient sources.

To demonstrate the emergent switch based metabolic strategies, based on the possibilities encoded in the genome, we consider phototrophic growth with two alternative sources of extracellular nitrogen. We assume that, in addition to the nitrogen source *N*_*x*_ considered above, there is a second source of extracellular nitrogen *N*_*y*_, whose uptake and conversion to the intracellular nitrogen precurser *N* is facilitated by a coarse-grained protein complex *T*_*Y*_. Compared to the complex *T*_*N*_, however, the synthesis of *T*_*Y*_ requires more amino acids and its catalytic turnover number *k*_cat_ is lower. Within their genome, strains may encode either of the two (coarse-grained) uptake protein complexes, *T*_*N*_ or *T*_*Y*_, or both. The respective strains are denoted as (*T*_*N*_)-strain, (*T*_*Y*_)-strain and (*T*_*N*_ +*T*_*Y*_)-strain. The inclusion of both protein complexes within the genome, however, entails additional cellular costs: a larger genome corresponds to a (slightly) higher fraction of the non-protein biomass *Q*. Moreover, additional protein machinery is required to facilitate cellular decision to control the expression of both enzyme complexes, resulting in an increased fraction of non-catalyzing proteins *P*_*Q*_. The parameterization of the strains is provided in the Materials and Methods.

In the following, we assume that the extracellular nitrogen source *N*_*y*_ is constantly available (analogous to, e.g., atmospheric dinitrogen), whereas the availability of the nitrogen source *N*_*x*_ varies. Figure 5 shows the growth curves of all three strains as a function of *N*_*x*_ in the presence of a basal availability of *N*_*y*_ (Figure 5A), as well as the expression of the respective uptake complexes for the (*T*_*N*_ +*T*_*Y*_)-strain (Figure 5B). As expected, the (*T*_*Y*_)-strain exhibits a constant growth rate, due to the constant basal availability of *N*_*Y*_. The (*T*_*N*_)-strain exhibits a Monod-type dependence on the availability of *N*_*x*_, as already shown in Figure 2. The combined (*T*_*N*_ +*T*_*Y*_)-strain, however, exhibits a switch between two growth regimes: for low availability of external *N*_*x*_, the strain expresses the protein complex *T*_*Y*_ and utilizes the nitrogen source *N*_*Y*_. In this regime, the (constant) specific growth rate is slightly below the rate observed for the (*T*_*Y*_)-strain due to the increased burden of non-catalytic biomass. If the availability of *N*_*x*_ exceeds a certain threshold, the (*T*_*N*_ +*T*_*Y*_)-strain switches its preferred nitrogen source and expresses the protein complex *T*_*N*_. The growth rate then increases with increasing availability of *N*_*x*_, though it always remains below the growth rate of the (*T*_*N*_)-strain (again due to the increased burden of non-catalytic biomass). Hence, we expect that the (*T*_*N*_ +*T*_*Y*_)-strain will be outcompeted in any constant environment, but will have a competitive advantage in (some) environments with variable nitrogen availability.

**Figure 5.**
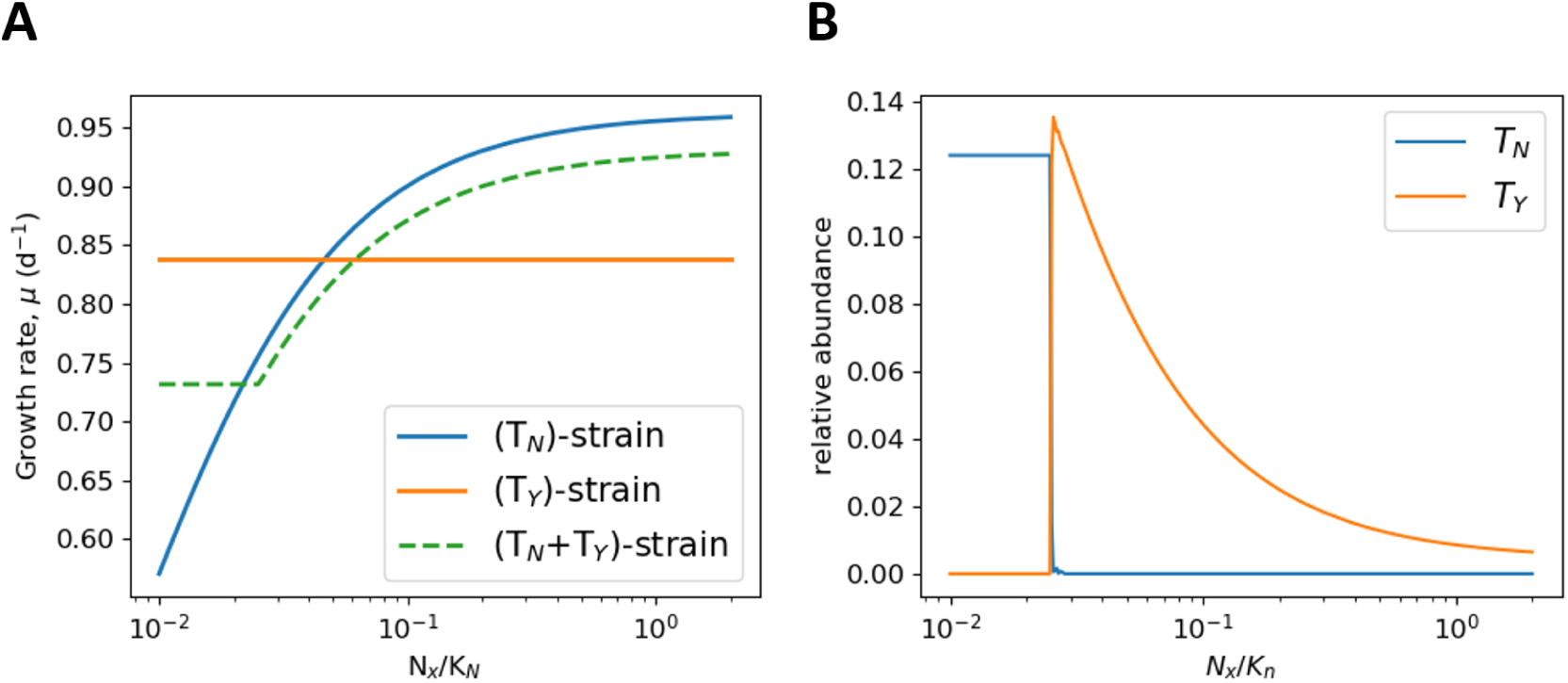
Panel A: The predicted specific growth rate of three different cyanobacterial strains at different concentrations of the external nitrogen source *N*_*x*_ and a basal supply of the alternative nitrogen source *N*_*y*_. The strains are denoted as (*T*_*N*_)-strain, (*T*_*Y*_)-strain and (*T*_*N*_ +*T*_*Y*_)-strain, and encode either a single uptake mechanism (*T*_*N*_ or *T*_*Y*_) or both within their genomes. The *T*_*N*_ +*T*_*Y*_)-strain has a higher biosynthesis cost in terms of increased genome size and additional regulatory proteins and hence exhibits a reduced specific growth rate compared to the streamlines strains. **Panel B:** Relative abundance (with respect to total proteome) of the nitrogen uptake mechanisms *T*_*N*_ and *T*_*Y*_ for the (*T*_*Y*_)-strain. The expression of the respective proteins depends on the environmental conditions.

For our purposes, the example serves to illustrate the following points: (i) biochemical resource allocation models build upon the genome of an organism and hence allow us to represent genetic diversity within strains, including genomes that encode several potential metabolic strategies and differences is genome size; (ii) the associated costs of larger genomes, including the the costs for additional expression of regulatory proteins can be incorporated into the parameterization of the model based on pan-genome-analysis and quantitative growth studies [31, 4, 63]; (iii) The optimal metabolic strategy for any given environment does not have to be specified in advance but is an emergent outcome of model simulation. Strains may switch between different strategies in different environments—with important consequences for ecosystem models; (iv) we observe that, within simulations, cells typically exhibit a hierarchy of preferred nutrients. That is, optimal solutions are not combinations of different uptake mechanisms. This behavior was previously proven for a different, but closely related, class of resource allocation models [62, 40], and is reminiscent of the phenomenon of catabolite repression. To what extent the hierarchy of preferred nutrients is a universal feature of microbial growth is insufficiently understood.

## A CASE STUDY: SEASONAL VARIATION AND CO-EXISTENCE

To exemplify the feasibility to utilize biochemical resource allocation models within ecosystems simulations, we consider a model of phytoplankton diversity recently proposed by Tsakalakis et al. [57]. We do not aim to recapitulate the full study of Tsakalakis et al. [57], but focus on the competition outcomes between opportunists (r-strategists) and gleaners (K-strategists) in constant versus time-varying environments. As shown above, the growth physiology of biochemical resource allocation models is an emergent property of the underlying biochemical parameters. We therefore assume that the biochemical parameters of carbon uptake, as well as nitrogen metabolism and uptake, differs between strains—reflecting the diversity of strains. As noted above, our premise is that enzyme-kinetic parameters are subject to physico-chemical trade-offs, for example trade-offs between the half-saturation constant and the maximal catalytic rate of an enzyme. We emphasize that such trade-offs are not an outcome of our modelling approach but need to be specified independently, for example based on detailed biochemical surveys and analysis [14]. We consider two strains of phytoplankton. The differences between both strains (detailed in Materials and Methods and Table 2) reflect trade-offs in the half-saturation constants and catalytic activities of the uptake mechanisms of nitrogen and carbon, and result in two functional groups of phytoplankton, gleaners and opportunists. The respective growth curves are shown in Figure 6. Gleaners are characterised by a higher affinity towards extracellular nitrogen, and an overall lower maximal growth rate. Opportunists are characterised by a high overall specific growth rate, but a lower affinity for extracellular nitrogen.

**Table 2.** Parameters of the model. The parameter values follow the data used in Faizi et al. [11]. If no data were available in the literature, the remaining parameters are estimated ^⋄^ based on generic values.

| Symbol | Definition | Gleaner/Control | Opportunist | Source |
| --- | --- | --- | --- | --- |
| $V_{cell}$ | Cell volume ($\mu m^3$) | 1.8 | 1.8 | [7] |
| $D$ | Rate of dilution ($d^{-1}$) | 0.25 | 0.25 | [10] |
| $d$ | Average cell density ($aa\ cell^{-1}$) | $1.4 \times 10^{10}$ | $1.4 \times 10^{10}$ | [11] |
| $k_d$ | Rate of photo damage | 0.56 | 0.56 | $\diamond$ |
| $\sigma$ | Effective absorption ($m^2\ \mu mol\ PSU^{-1}$) | 0.2 | 0.2 | $\diamond$ |
| $n_{PSU}$ | Size of photosynthetic unit $PSU$ ($aa\ molec^{-1}$) | 95451 | 95451 | [11] |
| $n_{TC}$ | Size of carbon transporter $T_c$ ($aa\ molec^{-1}$) | 1681 | 1681 | [11] |
| $n_{CB}$ | Size of Calvin-Benson (CB) proteins ($aa\ molec^{-1}$) | 2000 | 2000 | $\diamond$ |
| $n_c$ | Size of carbon metabolism ($M_c$) proteins ($aa\ molec^{-1}$) | 20000 | 20000 | $\diamond$ |
| $n_{TN}$ | Size of nitrogen transporter $T_n$ ($aa\ molec^{-1}$) | 10000 | 10000 | $\diamond$ |
| $n_P$ | Size of protein P ($aa\ molec^{-1}$) | 1000 | 1000 | $\diamond$ |
| $n_q$ | Size of metabolism protein $M_q$ ($aa\ molec^{-1}$) | 20000 | 20000 | $\diamond$ |
| $n_R$ | Size of ribosome $R$ ($aa\ molec^{-1}$) | 7358 | 7358 | [11] |
| $\gamma_{\max}$ | Maximal translation rate ($aa\ s^{-1}\ molec^{-1}$) | 22 | 22 | [6] |
| $K_C$ | Half-saturation constant of $T_c$ ($\mu M$) | 15 | 15 | [44] |
| $K_N$ | Half-saturation constant of $T_n$ ($\mu M$) | 10 | 50 | $\diamond$ |
| $k_{cat,PSU}$ | Turnover rate of $PSU$ ($s^{-1}$) | 250 | 250 | [37] |
| $k_{cat,TC}$ | Turnover rate of $T_c$ ($s^{-1}$) | 20 | 200 | $\diamond$ |
| $k_{cat,CB}$ | Turnover rate of $CB$ ($s^{-1}$) | 1 | 1 | $\diamond$ |
| $k_{cat,MC}$ | Turnover rate of $M_c$ ($s^{-1}$) | 10 | 10 | [37] |
| $k_{cat,TN}$ | Turnover rate of $T_n$ ($s^{-1}$) | 50 | 200 | $\diamond$ |
| $k_{cat,P}$ | Turnover rate of $P_q$ ($s^{-1}$) | 100 | 100 | $\diamond$ |
| $k_{cat,Q}$ | Turnover rate of $M_q$ ($s^{-1}$) | 100 | 100 | $\diamond$ |
| $Q$ | Relative abundance of $Q$ w.r.t. biomass | 0.5 | 0.5 | $\diamond$ |
| $P_Q$ | Relative abundance of $P_Q$ w.r.t. total proteome | 0.2 | 0.2 | $\diamond$ |

**Figure 6.**
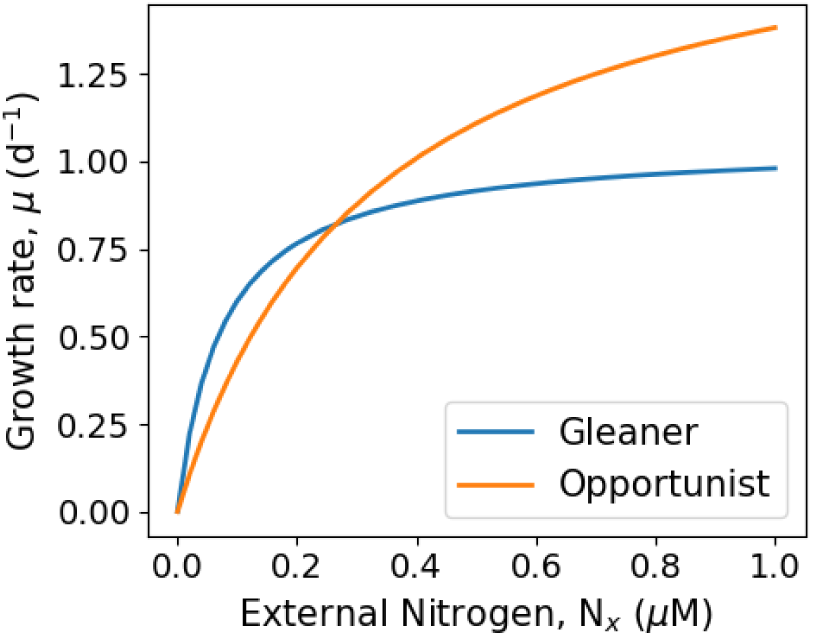
The growth curves of two competing strains of phytoplankton, opportunists and gleaners, in dependence of external nitrogen availability. The strains differ in the enzyme-kinetic parameters of their constituent enzyme complexes. Gleaners (K-strategists) have a growth advantage during phases of low external nitrogen availability, whereas opportunists (r-strategists) have a growth advantage at high concentrations of external nitrogen.

Following Tsakalakis et al. [57], we simulate the growth of both strains in two different environments: a constant light environment (control) and a light environment with seasonal variations in average light intensity. Extracellular inorganic carbon is assumed to be constant, a (single) source of extracellular nitrogen is is supplied via a constant influx. The dynamics of the abundances of gleaners (*ρ*_*G*_) and opportunists (*ρ*_*O*_) are described by the following ODEs

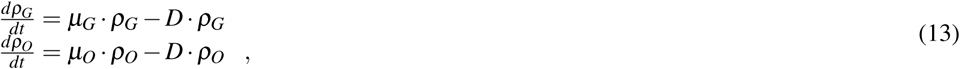

the dynamics of external nitrogen is described by

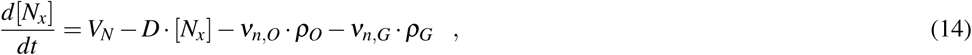

where *V*_*N*_ denotes a constant influx, and *ν*_*n,O*_ and *ν*_*n,G*_ denote the specific cellular uptake rates (as emerging from the respective models) of external nitrogen by the gleaners and opportunists, respectively. The population dynamics of both strains in constant and time-varying environments are shown in Figure 7. Simulations were performed using a Python ODE solver, the growth models are implemented a (series of) LP problems and solved at each time step. The procedure is computationally similar to dynamic FBA (dFBA), an established method for constraint-based analysis [34]. See Materials and Methods for details.

**Figure 7.**
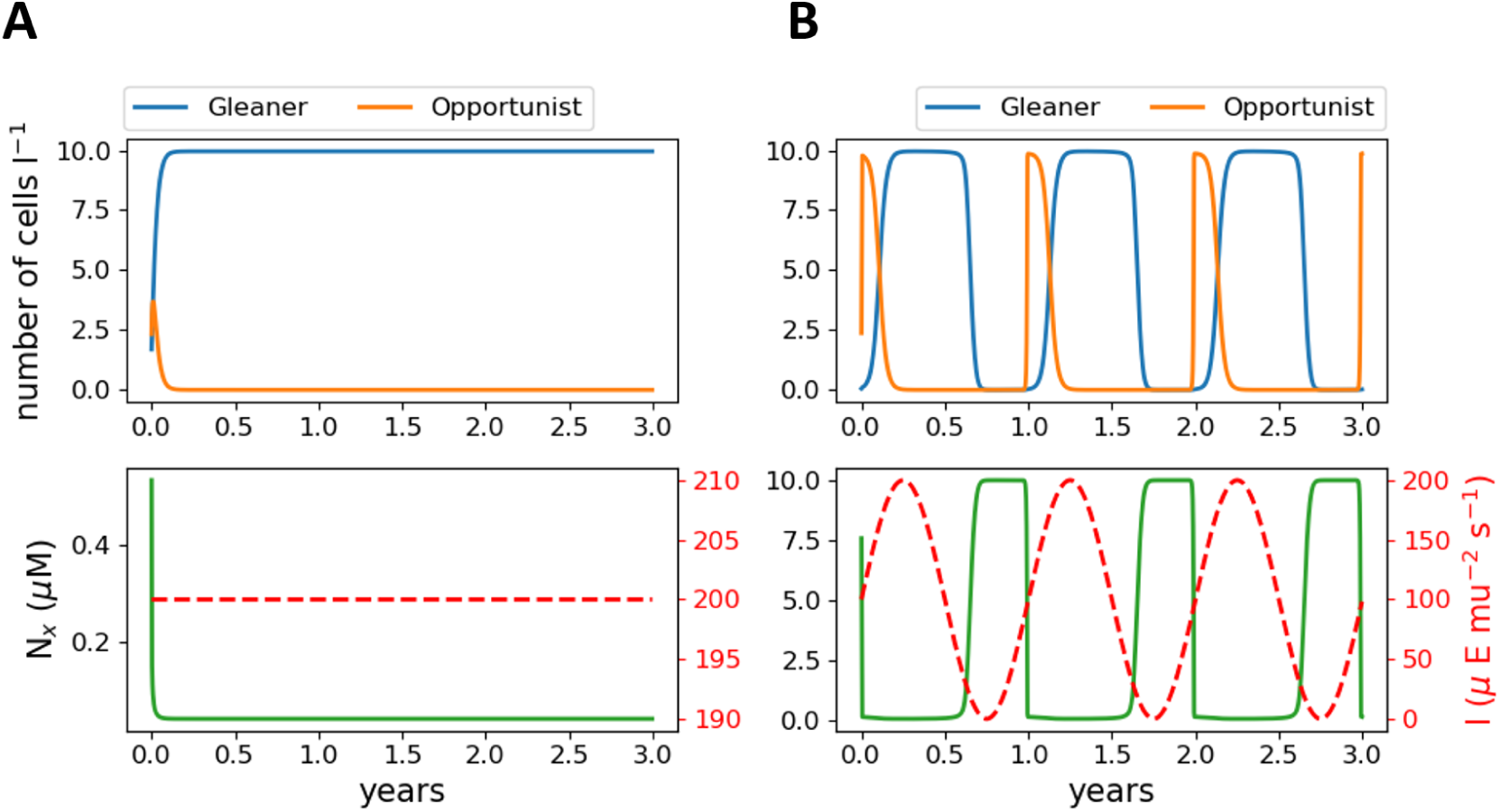
Population dynamics of opportunists and gleaners under different nutrient and light conditions. **Panel A:** shows competitive exclusions for constant light (*I* = 200*µEm*^*-*2^*s*^*-*1^), with the gleaner strain outcompeting the opportunist. **Panel B:** shows the co-existence of both strains. Whereas, the lower right panel shows the changes in the nitrogen concentration under the changing conditions of light intensities. All simulations are performed using the parameters given in Table 2.

As shown in Figure 7, gleaners outcompete opportunists in a constant light environment, consistent with the competitive exclusion principle. Seasonally changing light intensities, however, induce changes in strain abundances, and hence nitrogen availability. Temporal changes in nitrogen availability then result in the co-existence of both strains. During periods of low light availability, overall strain abundance decrease and the availability of extracellular nitrogen increases. With increasing light intensities, opportunists have a competitive advantage and quickly increase in abundance, thereby decreasing the availability of extracellular nitrogen and shifting the competitive advantage to gleaners until a decreasing light intensity restart the cycle. The simulation results are consistent the the corresponding simulations of Tsakalakis et al. [57] and demonstrate the feasibility to utilize biochemical resource allocation models in ecosystem simulations.

## DISCUSSION AND OUTLOOK

As emphasized by Follows and Dutkiewicz [15], there is currently a vast chasm between the ecologically and biogeochemically oriented parameterizations of growth utilized in ecological modelling and the metabolic-pathway perspective of microbial growth enabled by systems biology and modern genomics. The purpose of this study was to outline a connection between both fields and to show how recent biochemical models of microbial growth might contribute to close this chasm. In this respect, of particular interest are resource allocation models of microbial growth [18, 59, 49, 8, 11]. Biochemical resource allocation models allow us to provide a quantitative account of protein expression and biochemical processes based on knowledge about biochemical parameters. Our aim was to heed the call of Allen and Polimene [1] to provide growth models based on a robust physiological formulation that allow for trade-offs between resource allocation of competing physiological activities. We propose that biochemical resource allocation models, such as the ones described here, fulfill this paradigm towards a new generation of plankton models. While mechanistic growth models [54], resource-allocation and cost-benefit analysis [48, 17, 15, 30], as well as models based on optimality [46, 47], are well established in ecological modelling, the resource allocation models described here directly build upon the framework of metabolic network reconstruction and constraint-based analysis—and therefore reflect the advances in quantitative growth physiology enabled by systems biology and modern genomics. The predictions from biochemical resource allocation models are often in excellent agreement with detailed physiological studies of model strains [19, 63] making them a good starting point for the description of microbial growth.

Biochemical resource allocation models can be formulated for almost all microbial strains for which a reference genome is available. Supported by recent analysis of the cyanobacterial pan-genome [3, 55, 4], and the diversity of energy metabolism in microbes [13], we hypothesize that such models will follow a modular paradigm: there is only a limited number of fundamentally different metabolic strategies available and microbial organisms are a mix-and-match conglomerate of these strategies (with many combinations excluded for biophysical or energetic reasons). The enormous diversity of microbial metabolism then arises from further variations and adaptations of biochemical parameters (with possible trade-offs), as well as from differences in cellular resource allocation. For example, recent studies show that the observed significant differences in the maximal specific growth rates between genetically similar cyanobacterial strains are related linked to differences in resource allocation strategies (such as the amount of storage compounds or differences in the PSII/PSI ratio) [63]. We note, however, that variability and possible trade-offs in enzyme-kinetic parameters are not an intrinsic part of biochemical resource allocation models but have to be provided as externallt—based on detailed biochemical studies [14]. The analysis of biochemical resource allocation models therefore distinguishes between trade-offs that arise from physicochemical constraints in enzyme evolution and trade-offs that arise from differences in protein expression and resource allocation.

The merits of biochemical resource allocation models are as follows: (I) the models can be formulated using different levels of complexity, from genome-scale representations taking into account all individual enzymes [19, 49], to intermediate representations [50], to coarse-grained models that consider protein complexes corresponding to (agglomerated) cellular processes, such as the model outlined above; (II) model parameterization is be based on available knowledge provided in biochemical databases [26] and our increasing knowledge about quantitative cell physiology [37]. The models therefore provide a link between physicochemical constraints of enzyme-kinetic parameters and observed growth kinetics. Key parameters for model parameterization are enzyme costs (in terms of amino-acids and co-factor requirements) and enzymatic catalytic activities. Information about regulatory mechanisms is not required; (III) the models allow to represent metabolic diversity by taking distributions of parameters (and possible trade-offs) into account. Biochemical resource allocation models therefore allow to implement selection-based approaches [15]—following the Baas-Becking paradigm “everything is everywhere but environment selects” (cited after Follows and Dutkiewicz [15]); (IV) biochemical resource allocation models allow for complex metabolic behavior, such as switches between different metabolic strategies. Most microbes are capable of more than one metabolic mode and conventional Monod-type models face difficulties to describe transitions between metabolic modes. For biochemical resource allocation models the modes of energy generation or nutrient uptake strategies (and hierarchies) emerge without further specification as part of the optimization procedure. (V) the latter also allows model to be embedded within evolutionary simulation to explain how different metabolic strategies and strains with different genome sizes may emerge and co-exist. (VI) biochemical resource allocation models of the form discussed here only require linear optimization and hence are computationally tractable. While it is (currently) not possible to formulate kinetic models at the genome-scale, the implementation of bioechemical resource allocation models is computationally feasible even for large models [19, 49]. Coarse-grained models, such as the one discussed above, can be solved fast and efficiently and hence are suitable for ecosystems simulations. In case computational capacity is limiting, it is possible to devise approximate schemes (such as lookup tables).

Notwithstanding their merits, current biochemical resource allocation models are not (yet) the panacea for ecological simulations. We expect that different approaches are needed, as well as further improvements of biochemical resource allocation models and other whole-cell systems biology models. In particular, current biochemical resource allocation models can be extended along the following lines: (I) current simulations typically focus on steady state analysis. While it has been shown, that biochemical resource allocation models can be solved for time-varying environments [50, 49], the computational burden is significant. Nonetheless, it is paramount importance to be able to represent phenomena such as storage, bet-hedging or luxury uptake of scarce nutrients (i.e, the uptake of nutrient beyond what is currently required in anticipation of possible future limitations). These phenomena are also aspects of resource allocation strategies and hence can be represented by appropriate models. (II) currently models are based on a metabolic perspective of growth. In principle, also trade-offs between growth and other cellular properties can be considered, such as the resilience against stress or predation. (III) a better understanding of physicochemical trade-offs in enzyme-kinetic parameters is required. Further quantitative growth studies, along the lines of Zavřel et al. [63] are required to quantify the cost of regulation for strains with different genome sizes.

Overall, we envision a unified framework to construct biochemical resource allocation models based on reference genomes and suitable biochemical parameterizations. Such models will allow us to represent the microbial diversity observed in almost all environments and will open up new avenues to interface biogeochemical and ecological questions with recent knowledge obtained from quantitative microbial growth physiology.

## MATERIALS AND METHODS

### Biochemical resource allocation models

The implementation of the resource allocation models follows the algorithms described in [18, 50, 49]. A model consists of two types of components: steady-state metabolites and cellular macromolecules (which include catalytic protein complexes and quota components). We assume that the internal metabolites are at a quasi-steady state, *i*.*e*., metabolism readjustments are faster than changes in the external environmental. Thus, the concentrations of internal metabolites are not explicitly evaluated in the model, and the metabolic network is assumed to be balanced at all times. We neglect dilution by growth of internal metabolites. The quota components (protein *P*_*Q*_ and remaining biomass *Q*) fulfill no explicit function within our model and their synthesis is enforced using fixed quotas (except otherwise noted, the quota protein component *P*_*Q*_ is assumed to be 20% of total protein, the non-protein biomass *Q* is assumed to be 50% of total biomass).

### The biochemical resource allocation model of phototrophic growth

The biochemical resource allocation model shown in Figure 1 is assembled following the stoichiometry and data described by Faizi et al. [11]. We note that the model of Faizi et al. [11] is a nonlinear kinetic ODE model, hence computationally different from the model described here. Growth is facilitated by 8 protein complexes: 6 enzyme and transport complexes, ribosomes *R* and a non-catalyzing quota protein compenent *P*_*Q*_. The enzyme and transporter catalyze the following reactions:

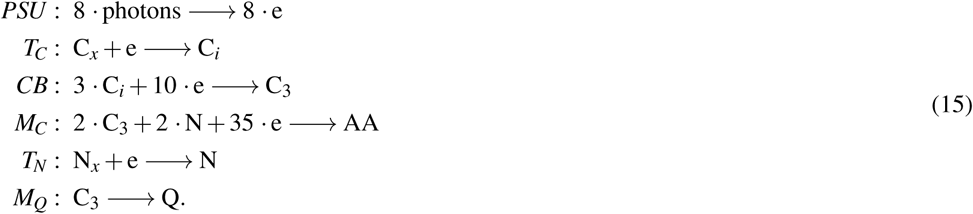

Protein translation is described by the equation

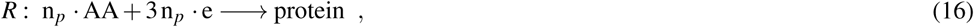

where *n*_*p*_ denotes the size of the respective protein in amino acids.

### Capacity constraints of catalytic enzymes

All enzyme-catalyzed reactions are constrained by the amount of the respective enzyme, according to equation (7). The constraints for uptake and light harvesting reactions also depend on the availability of the respective substrates. In particular, for (i) the uptake of inorganic carbon,

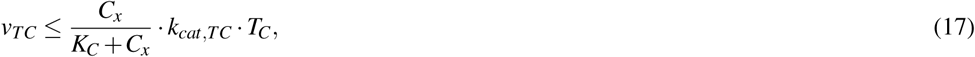

(ii) for uptake of extracellular nitrogen,

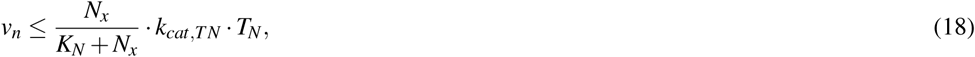

and (iii) and for light harvesting and photosynthesis

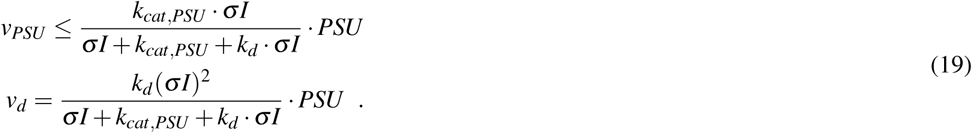

The equation for light harvesting and photosynthesis is derived from a two-state model of photosynthesis that accounts for photodamage, see [11] for a derivation.

The constraints on the ribosomal capacity are

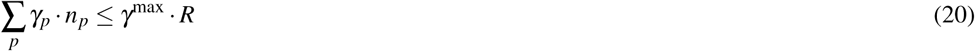

where *n*_*p*_ denotes the protein size (in amino acids per molecule), *γ*_*p*_ its translation rate, and *γ*^max^ denotes the maximal translation rate of ribosomes. We note that all capacity constraints can be implemented as linear constraints.

### Solving the resource allocation model as a LP

For any given set of external parameters *C*_*x*_, *N*_*x*_, *I* and specific growth rate *µ*, the model implemented as a linear program *LP*(*µ*). The problem is described by three matrices ***N***, ***B***, and ***C***, the vector of reaction rates ***v*** = (*v*_*i*_, *γ*_*k*_)*T* (including metabolic and translation rates), and the vector ***P*** of macromolecules. The constraints are

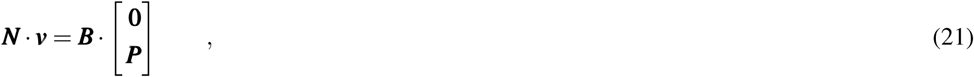

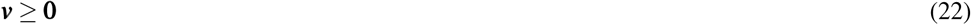

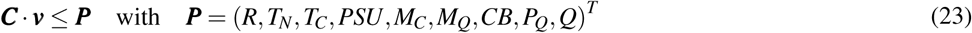

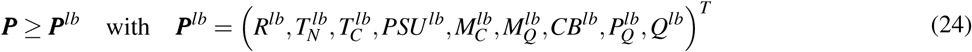

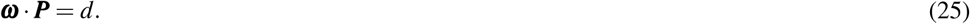

Constraint (21) enforces mass-balance at steady-state, including terms for dilution (with dilution of metabolites neglected). The matrix ***B*** is a diagonal matrix with elements *µ* on the diagonal. Constraint (23) described the (linear) enzymatic capacity constraints, the matrix ***C*** is largely diagonal, except for the constraints on the translation rate. Constraint (24) provides a lower bound for the abundance of each macromolecules (which is zero except for the quota components). Constraint (22) ensures positive fluxes in the LP problem. Constraint (25) enforces a constant cell density. The vector ***ω*** described the sizes of the macromolecules (in units of amino acids per molecule) and *d* denotes the cell density (in units of amino acids per cell).

The above described LP is solved as a feasibility problem for a given *µ*. To obtain a solution for the maximal specific growth rate in a given environment, the global optimum of *µ* is found using bisection, analogous to the method used in [50, 49].

### Model parameterization

A complete list of model parameters is provided in Table 2. Parameterization follow the data used in Faizi et al. [11]. The size of macromolecules is estimated using the size of an average enzyme times the approximate number of steps used in the pathway. The size of the protein complex *P*_*Q*_ and the biomass component *Q* is arbitrary. Turnover rates are chosen according to average values described in [37]. The description of the photosystem is adopted from Faizi et al. [11], with *σ* denoting the effective absorption cross-section per photosystems, and *k*_*d*_ the rate of photodamage.

To simulate the growth on two alternative sources of external nitrogen, we used the set of additional parameters given in Table 3.

**Table 3.**
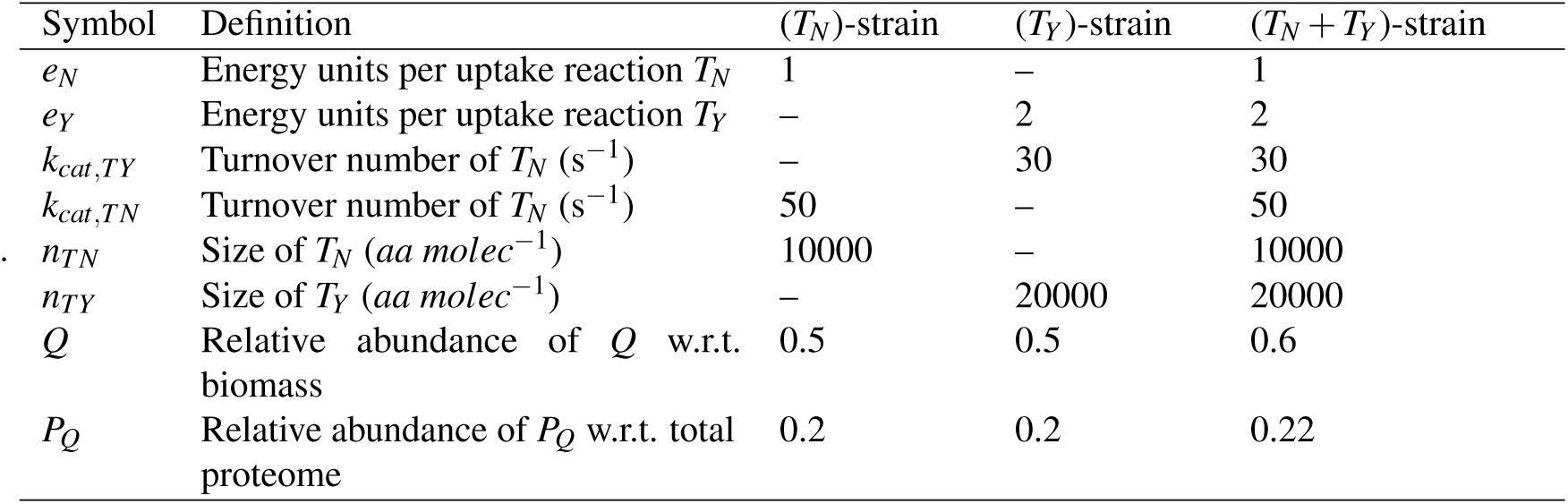
Specific parameters used to model growth on two alternative sources of external nitrogen. The remaining parameters are same as described in Table 2

| Symbol | Definition | ($T_N$)-strain | ($T_Y$)-strain | ($T_N + T_Y$)-strain |
| --- | --- | --- | --- | --- |
| $e_N$ | Energy units per uptake reaction $T_N$ | 1 | – | 1 |
| $e_Y$ | Energy units per uptake reaction $T_Y$ | – | 2 | 2 |
| $k_{cat, TY}$ | Turnover number of $T_N$ ($s^{-1}$) | – | 30 | 30 |
| $k_{cat, TN}$ | Turnover number of $T_N$ ($s^{-1}$) | 50 | – | 50 |
| $n_{TN}$ | Size of $T_N$ ($aa \text{ molec}^{-1}$) | 10000 | – | 10000 |
| $n_{TY}$ | Size of $T_Y$ ($aa \text{ molec}^{-1}$) | – | 20000 | 20000 |
| $Q$ | Relative abundance of $Q$ w.r.t. biomass | 0.5 | 0.5 | 0.6 |
| $P_Q$ | Relative abundance of $P_Q$ w.r.t. total proteome | 0.2 | 0.2 | 0.22 |

### The computational modelling framework

A computational model is developed using Python as a programming language. The framework uses functionality from the following packages: numpy [43], scipy [27], matplotlib [25], pandas [36], sundials [24] and Gurobi [20]. In particular, we use Gurobi for solving the LP-based optimisation and CVODE integrator from the sundials package to solve the system of ODEs. The version of the modelling framework used to produce the results presented in this manuscript is publicly available with instructions to install and run simulations at (https://github.com/surajsept/cyanoRBA). The development version is hosted on GitLab (https://gitlab.com/surajsept/RBmodels) and people interested in contributing can request access by contacting the author (S.S.).

